# DHA but not EPA induces the trans-differentiation of C2C12 cells into white-like adipocytes phenotype

**DOI:** 10.1101/2021.03.19.436137

**Authors:** Saeed Ghnaimawi, Lisa Rebello, Jamie Baum, Yan Huang

**Affiliations:** Department of Cell and Molecular Biology Program, University of Arkansas, Fayetteville, USA; Department of Food Science, Division of Agriculture, University of Arkansas, Fayetteville AR, USA; Department of Animal Science, Division of Agriculture, University of Arkansas, Fayetteville, USA

**Author notes:** Correspondence to: Dr. Yan Huang, Department of Animal Science, Division of Agriculture, University of Arkansas, Fayetteville, USA, University of Arkansas, Fayetteville, AR 72701. **Conflict of interest statement**. The authors declare that the research was conducted in the absence of any commercial or financial relationships that could be construed as a potential conflict of interest. **Author contribution statement**. The experiments was designed by Yan Huang, Saeed Ghnaimawi, and Jamie Baum. Saeed Ghnaimawi performed the research, analyzed the data, and wrote the paper. Lisa Rebello contributed the training of seahorse analytic tool and the protocol troubleshooting.

**Keywords:** N-3 PUFAs, C2C12 cells, brown adipogenesis, white adipogenesis, myogenesis, Trans-diffrentiation

## Abstract

Muscle derived stem cells (MDSCs) and myoblast play an important role in myotube regeneration when muscle tissue is injured. However, these cells can be induced to differentiate into adipocytes once exposed to PPARγ activator like EPA and DHA that are highly suggested during pregnancy. The objective of this study aims at determining the identity of trans-differentiated cells by exploring the effect of EPA and DHA on C2C12 undergoing differentiation into brown and white adipocytes. DHA but not EPA committed C2C12 cells reprograming into white like adipocyte phenotype. Also, DHA promoted the expression of lipolysis regulating genes but had no effect on genes regulating β-oxidation referring to its implication in lipid re-esterification. Furthermore, DHA impaired C2C12 cells differentiation into brown adipocytes through reducing the thermogenic capacity and mitochondrial biogenesis of derived cells independent of UCP1. Accordingly, DHA treated groups showed an increased accumulation of lipid droplets and suppressed mitochondrial maximal respiration and spare respiratory capacity. EPA, on the other hand, reduced myogenesis regulating genes, but no significant differences were observed in the expression of adipogenesis key genes. Likewise, EPA suppressed the expression of WAT signature genes indicating that EPA and DHA have an independent role on white adipogeneis. Unlike DHA treatment, EPA supplementation had no effect on the differential of C2C12 cells into brown adipocytes. In conclusion, DHA is a potent adipogenic and lipogenic factor that can change the metabolic profile of muscle cells by increasing myocellular fat.

## 1. Introduction

The skeletal muscle tissue making up the majority of body mass is manifested by its responsibility of body movement and metabolism regulation. Muscle fibers are generated prenatally during fetal development while they increase in size and length during the postnatal stage (1). Prenatally, somite-derived myoblasts, the progenitors of muscle tissue, leave the cell cycle after proliferation to form myotubes in a process known as myogenesis (2). Progenitor cells, including muscle-derived stem cells (MDSCs) and myoblasts play an important role in regenerating muscle fibers following injury. The cells responsible for the repairing process are normally, but gain a massive capacity of rapid proliferation and differentiation to form new myofibers (3).However, these cells can also be induced to differentiate into other lineages such as adipocytes, osteocytes, and chondrocytes in response to external stimuli like maternal nutrition (4).

Harper and Pethick (5) emphasized that intramuscular fat (IMF) can be originated from mesodermal derived multipotent stem cells, which are a regular source of adipocytes, and from myoblasts and fibroblasts upon exposure to adipogenic factors. Hence, there is an urgent necessity to identify stimuli that are capable of committing myoblasts into adipocyte lineage and to discover the molecular mechanisms orchestrating the trans-differentiation process. Such studies may help find out more about the origin of intramuscular adipocytes, whether it is ectopic extension adipocytes from other fat depots or by-products of possible trans-differentiation of myoblasts into adipocytes.

Health benefits of long-chain (n-3) polyunsaturated fatty acids (PUFAs) supplements, particularly eicosapentaenoic acid (EPA) (20:5n3) and docosahexaenoic acid (DHA), have been well established (6–9). Several studies have attested that EPA and DHA have an important role in blunting muscle atrophy, orchestrating lipid metabolism, enhancing protein synthesis, and improving insulin resistance in skeletal muscle (10–12). It was stressed that 30 µM DHA is sufficient to counteract the effect of palmitate-induced insulin resistance in C2C12 myotubes (11). Furthermore, Kamolrat, et al. (10) found that muscle protein synthesis can be considerably improved after exposing to 50 µM EPA. However, the effect of EPA and DHA on the myogenesis process and the possibility of reprogramming myoblasts into another lineage is still controversial. Consequently, further investigations are required to determine the association between the usage of EPA and DHA and myoblast fate (12).

N-3-PUFA supplementations, particularly EPA and DHA, are highly recommended during pregnancy because of their beneficial effects on fetal neuronal development along with improving retinal and immune functions (13). However, long-term exposure to n-3 PUFAs carries the risk of growth deficiencies that might be linked to nutritional toxicity (14). Considering the potential impact of maternal nutrition on myoblast proliferation and differentiation during fetal development (15), we hypothesized that EPA and DHA supplementation may induce myoblast trans-differentiation into adipocyte lineage while compromising fetal skeletal muscle development, generating long-term impacts on insulin sensitivity and energy balance of muscle in progeny.

It has been highlighted that the adipogenesis process can be induced in non-adipogenic cells, such as fibroblasts and myoblasts, by enforcing the ectopic expression of PPARγ, the key gene in regulating the adipogenic differentiation (16, 17). Anticipating the potential activation of PPARs by n-3 polyunsaturated fatty acids, well-known PPARs’ ligands (18), there is a great chance that the maternal diet enriched with EPA and DHA may induce trans-differentiation of myoblasts into adipocytes. As reported by (18), n-3 PUFA enriched diet is positively correlated with increasing the accumulation of intramuscular fat where the adipogenic effect was attributed to over-expression of adipogenesis signature genes. Furthermore, EPA has been recently identified to promote the conversion of myoblasts cells into the adipogenic lineage (19). However, the identity of the adipocytes derived from myoblasts is still needed to be defined. Therefore, the objective of this study was to explore the effect of EPA and DHA on myoblasts undergoing differentiation into brown and white adipocytes in an *in vitro* model. Considering the different functions of brown and white adipose tissue, potential expansion of BAT mass can enhance fatty acid oxidation, insulin sensitivity, and thermogenesis holding promise to encounter some pathophysiological disorders such as dyslipidemia and obesity. Conversely, WAT expansion may enhance the incidence of insulin resistance, lipotoxicity, and obesity. In our previous study we found that the combined supplementation of EPA and DHA in culture medium induced C2C12 cells reprograming into white adipocyte-like phenotype (20, 21). Our primary focus here was to determine the participation level of each one of these n-3 polyunsaturated fatty acids in the adipogenesis process. Accordingly, the investigation was carried out to find the independent roles of EPA and DHA on the potential differentiation of brown or white adipocytes from C2C12 cells.

## 2. Materials and methods

### 2.1. **Cell culture and differentiation**

C2C12 cell culture was conducted as previously described by Klemm, et al. (22). Briefly, the cells were initially cultured in a growth medium composed of Dulbecco’s modified Eagle’s medium (DMEM) 89% supplemented with 10% fetal bovine serum and 1% penicillin-streptomycin (Sigma-Aldrich, St. Louis, Missouri United States) at 37 °C in a 5% CO_2_ atmosphere. Confluent cells were then treated with a white (WIDM) and brown adipogenic differentiation induction medium (BDIM) in the absence (CON) or presence of doses of 50µM EPA or DHA for 10 and 7 days, respectively. White adipogenic differentiation induction medium 1 (WDIM 1) was composed of 0.125 mM IBMX, 0.3 µM dexamethasone, 1µg/ml insulin for the first 6 days. Then, the cells were switched into WDIM 2 composed only of 1µg/ml insulin for 4 days (23). On the other hand, browning differentiation induction medium (BDIM 1) was consisted of 0.1 mM IBMX, 125 nM indomethacin, 1 µM dexamethasone, 5µg/ml insulin, 1 nM T3 and 5 µM rosiglitazone. After 3 days the cells were exposed to BDIM 2 composed of 5µg/ml insulin, 1 nM T3 and 5 µM rosiglitazone (24). All the reagents composing WDIM and BDIM are from (Sigma-Aldrich, St. Louis, Missouri United States).

### 2.2. Oil Red O (ORO) staining

At the last day of differentiation in both experiments, the medium was aspirated off and the cells were rinsed three times in Phosphate Buffer Solution (PBS). Cells were fixed with 10% neutral buffered paraformaldehyde (Sigma-Aldrich, St. Louis, Missouri United States) for 30 minutes at room temperature followed by 3 times washing with PBS. Lipid droplets were stained with ORO (Sigma-Aldrich, St. Louis, Missouri United States) (25). A 0.5g of ORO was dissolved in 100 ml absolute isopropanol to prepare stocking solution. Six parts of stocking solution was mixed with four parts of distilled water to make working solution and then filtered using 0.2μm filter. Cells were maintained in ORO working solution for 30 minutes at room temperature 22^°^ . Morphological changes were observed under the microscope and intracellular lipid droplets were identified in the cytoplasm by their bright red color. Nikon DS-Fi3 digital camera mounted on a Nikon Eclipse TS 2R light microscope (Konan, Minato-ku, Tokyo, Japan) was used for imaging and the results were analyzed using ImageJ software. Finally, ORO was extracted by 1 ml 100% (v/v) isopropyl alcohol to quantitatively measure the density of the stain in different groups at 520 nm using microplate reader (Gene 5 Bio Tek Spectrometer, BioTek, northern Vermont, USA). Isopropanol was used as a blank (26)

### 2.3. Real-time PCR

Total RNA was extracted from the cells using RNeasy mini Kit (QIAGEN, Germany). The cDNA was synthesized using Χ iScript kit (Bio-Rad, Richmond, California) by following the manufacturer’s protocol. Real-time PCR was performed using CFX Connect Real-Time PCR Detection System (Bio-Rad, Richmond, California). Primers (Table 1) used were designed in accordance with NCBI database and Primer Quest (IDT. com). The data were normalized to the reference gene 18s, and the relative expression levels were expressed as fold change (27).

**Table 1:**
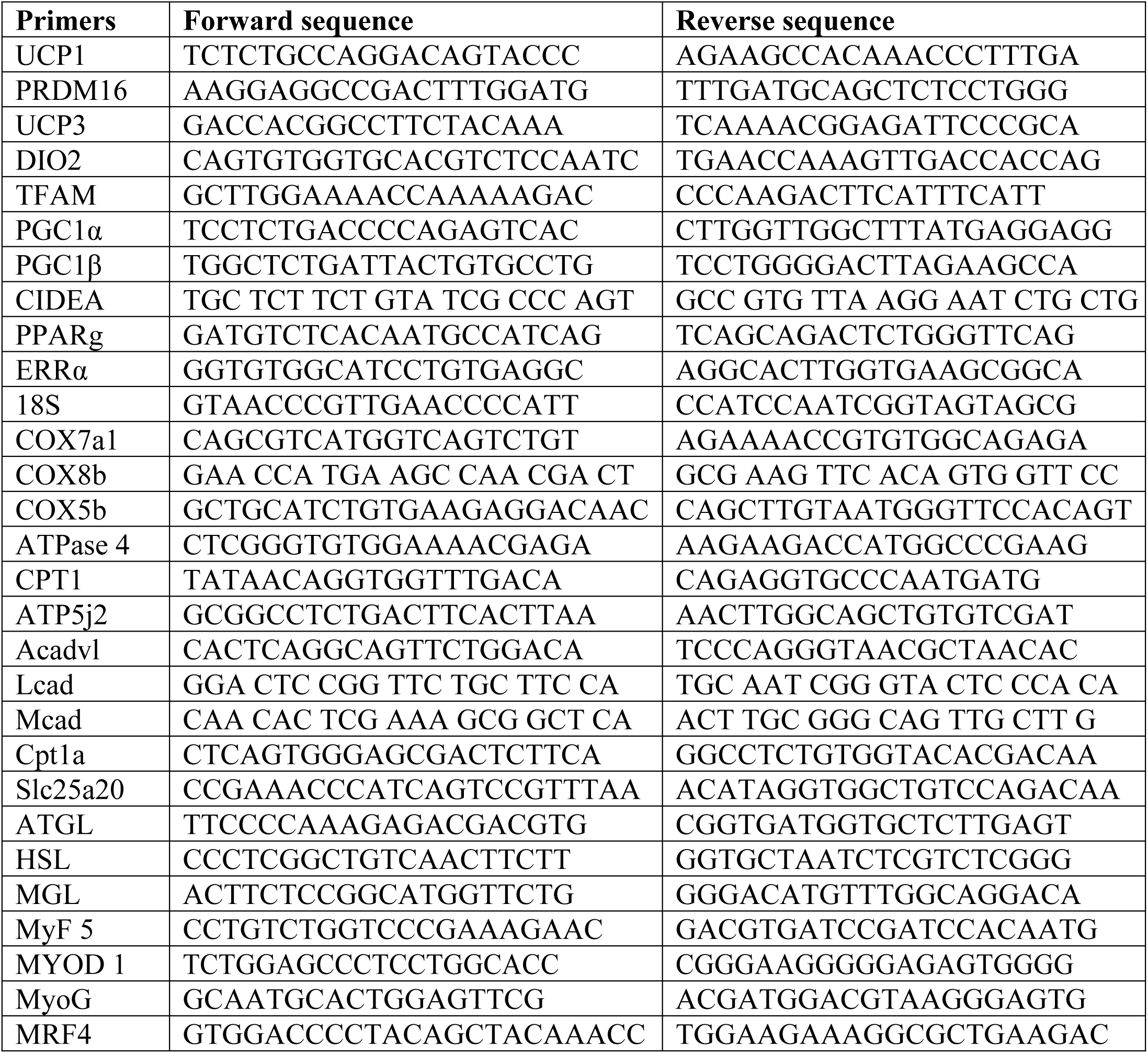
Primer sequences for real-time PCR

### 2.4. Oxygen consumption rate (OCR)

Mitochondrial function was evaluated to assess typical bio-energetic profile of cells by directly measuring Oxygen consumption rate (OCR) in differentiated groups using Seahorse XFP (Seahorse Bioscience, Santa Clara, United States, www.seahorsebio.com) and Agilent Seahorse XFP Cell Mito Stress Test (Seahorse Bioscience, Santa Clara, United States). Cells were seeded in customized Seahorse 8-well plates at an initial density of 10× 10^3 cell/ well and induced to differentiate into brown and white adipocytes as described above. One day before the assay, the cartridge was hydrated and incubated overnight in non CO2 incubator at 37 C. On the day of assessment, cells were incubated for 1 hour with completed assay medium (PH 7.4) consisted of basal medium supplemented with 10 mM glucose, 10mM pyruvate, and 2 mM glutamine at 37 °C in a non-CO2 incubator. During the assay, the following chemicals including: oligomycin, FCCP, and rotenone/ antimycin-A, were orderly injected at final concentrations of 2μM, 0.7 μM, and 1 μM respectively (All from Seahorse Bioscience). The Seahorse XF Cell Mito Stress Test Report Generator was used to analyze the data. The results were normalized to protein concentration (28).

### 2.5. Western blot assay

Cells were isolated using PBS (1 ml/well), and protein, then, was extracted by lysis buffer, T-PER, (Sigma-Aldrich, St. Louis, Missouri United States) mixed with protease and phosphatase cocktail inhibiter at a ratio of 100:1. Protein concentration was measured using Pierce™ BCA Protein Assay Kit (Sigma-Aldrich, St. Louis, Missouri United States) following the manufacturer’s instructions. Samples were separated on Mini-PROTEAN precast gels (Bio-Rad) and transferred onto Trans-Blot ® Turbo™ Mini PVDF Transfer Packs (Bio-Rad). Immuno-staining was conducted by overnight incubation with primary antibodies [GAPDH (1:1000, Cusobio, Waltham, MA, USA), myogenic differentiation 1 (MyoD) (1:25000, Cusobio), MyoG (1:25000, Cusobio), UCP1 (1/1,000; Abcam), UQCRC1 (complex III) (1/1,000; Invitrogen, Waltham, Massachusetts, USA), SDHA (complex II) (1/1,000; Invitrogen), PRDM16 (1/1,000; Santa Cruz).and PGC1α (1:1000, Abcam), C/EBPβ (1:1000, Cusobio), PPARγ (1:1000, cell signaling), Bmp4(1:1000, Cusobio), Myogenic factor 6 (MRF4) (1:1000, Cusobio)] . The membrane was then incubated with HPR conjugated secondary antibody for 1 hour after 3 times washing with TBST buffer. The bands were visualized using ECL immunoblotting clarity system (Bio-Rad, California, USA) and detected on ChemiDoc TM Touch imaging system (Bio-Rad). Band density was normalized according to the glyceraldehyde-3-phosphate dehydrogenase (GAPDH) content and analyzed using Image Lab software (Bio-Rad, California, USA) (29).

### 2.6. Statistical analysis

All data from assays used to compare CON with EPA or DHA treated group were assessed for significance by the unpaired Student’s t-test and one-way ANOVA analysis of variance, and arithmetic means ± SEM are reported to express the results. *P<*0.05 was considered statistically significant.

## 3. Results

### 3.1. Effect of EPA and DHA separately on white adipogenesis of C2C12 cells

#### 3.1.1. Both EPA and DHA suppressed the expression of genes regulating the differentiation of C2C12 cells in to mature myotubes

Exposing to EPA or DHA respectively suppressed myotubes formation in C2C12 cells exposed to EPA and DHA separately. Both EPA and EPA down-regulated the expression levels of *MyoD1*, *MyoG*, and *MRF4* but not *Myf5* indicating that the myogenesis process was compromised during the early stage of myogenic differentiation. DHA treatment decreased the expression levels of *MyoD1*, *MyoG*, and *MRF4* by (74% ± 3.9, 57% ± 21.7, and 97% ± 0.36 respectively). On the other hand, the expression levels of *MyoD1*, *MyoG*, and *MRF4* were down-regulated in EPA treated cells by (49% ±9.1, 89%± 4.5, and 93% ±0.63 respectively) **(Figure 1).**

**Figure 1:**
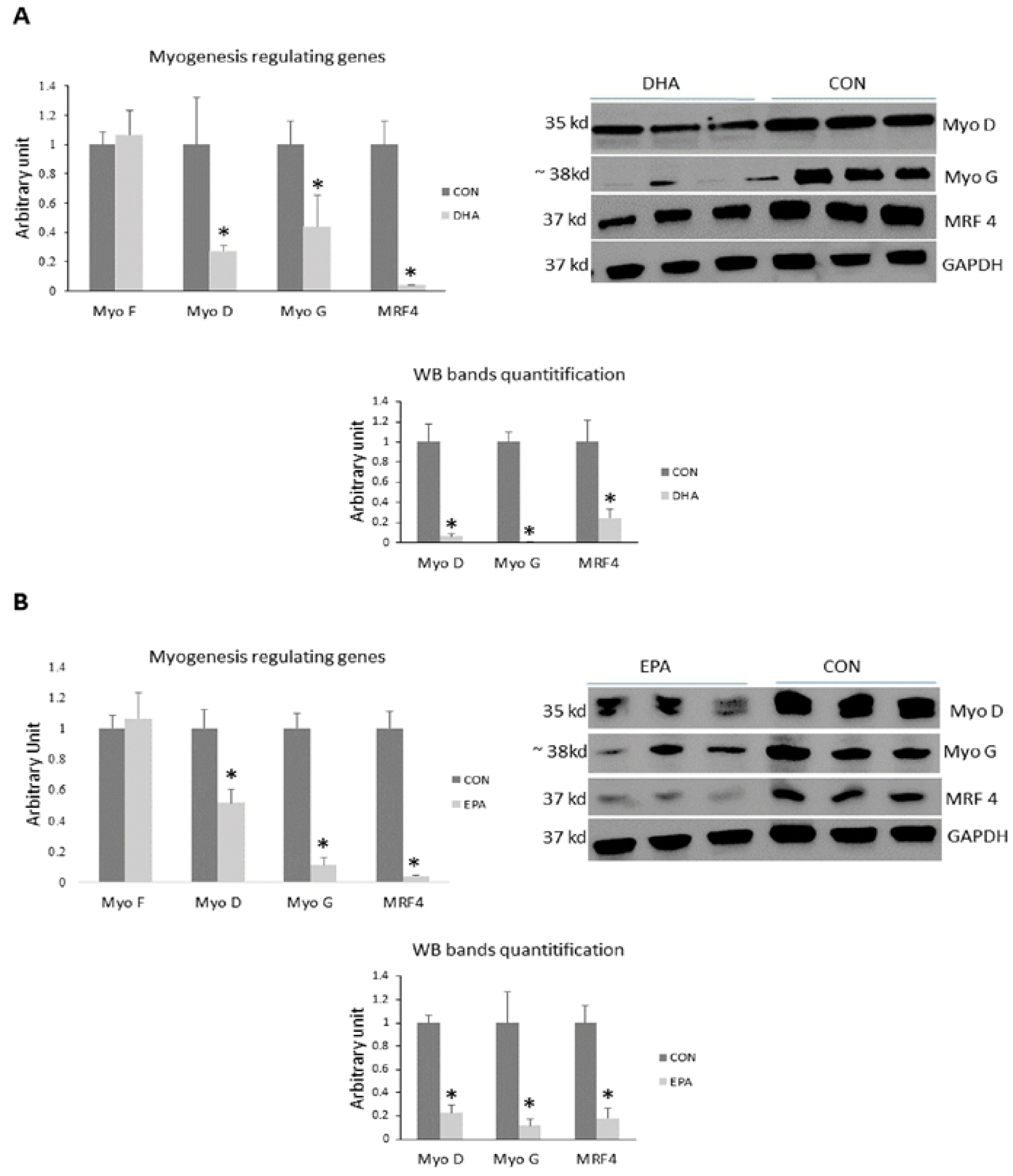
Representative image of western blot, densitometric analysis, and quantitative RT-qPCR analysis of the expression levels of myogenesis regulating genes in C2C12 cells during the trans-differentiation into white adipocytes by a hormonal cocktail. The cells were treated with differentiation induction medium (DIM) in the absence (CON) or presence of individual doses of 50 µM EPA and 50 µM DHA for 10 days. All the values of RT-qPCR were normalized to the expression levels of 18s genes and calculated in arbitrary units, n = 6. Data are represented as mean ± SEM. The significant differences is presented as *P < 0.05, Student’s t test. The level of GAPDH was used as a housekeeping gene to normalize the Western blots data. (A) Effect of DHA on the expression of myogenesis regulating genes. (B) Effect of EPA on the expression of myogenesis regulating genes.

#### 3.1.2. DHA but not EPA induced trans-differentiation of C2C12 cells into white adipocyte-like phenotype through up-regulation of WAT specific markers

We investigated the effect of isolated EPA and DHA on the expression levels of transcription factors regulating white adipogenesis such as *PPARγ, C/EBPα*, *C/EBPβ*, *AP2*, *FAT*, *Bmp4*, *and Zfp423*. The expression levels of *PPARγ*, *C/EBPα, C/EBPβ, AP2, FAT, Bmp4,* and *Zfp423* in DHA treated cells were increased by (150% ± 9.5, 100% ± 25, 41%± 12.1, 628% ± 87.9, 249% ± 45.8 P= 0.034, 48% ± 9.3, and 84%± 29.2, respectively). On the other hand, EPA did not affect in the expression levels of basal adipogenesis key genes: *PPARγ*, *C/EBPα*, *C/EBPβ*, *AP2*, and *FAT*. However, EPA dramatically reduced the expression of WAT regulators including *Bmp4* and *Zfp234* by (745 % ±32.6 and 67% ± 2.9 respectively) when compared to control **(Figure 2)**.

**Figure 2:**
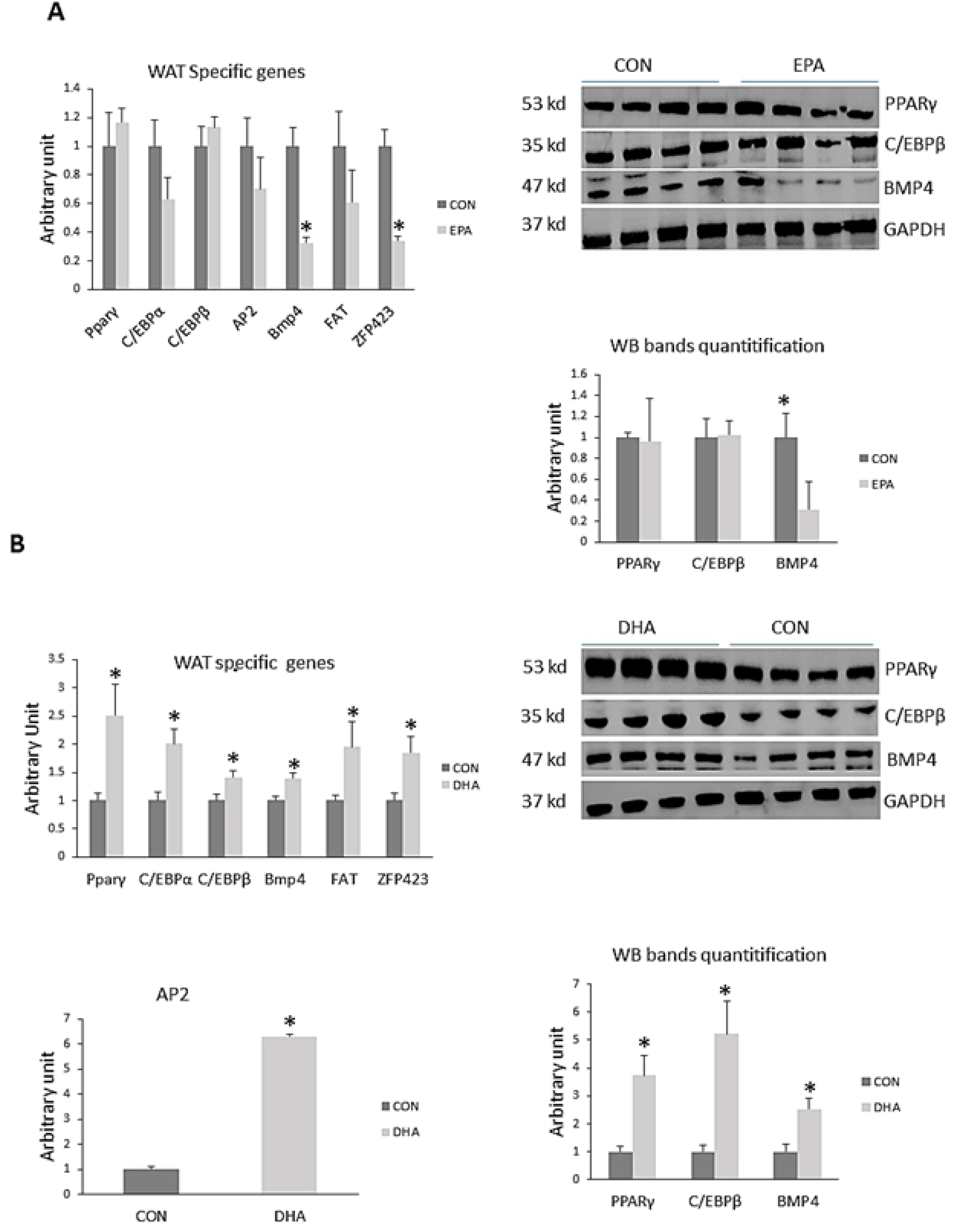
Representative image of western blot, densitometric analysis, and quantitative RT-qPCR analysis of the expression levels of adipogenesis regulating genes in C2C12 cells during the trans-differentiation into white adipocytes by a hormonal cocktail. The cells are treated with differentiation induction medium (DIM) in the absence (CON) or presence of individual doses of 50 µM EPA and 50 µM DHA for 10 days. All the values of RT-qPCR are normalized to the expression levels of 18s genes and calculated in arbitrary units, n = 6. Data are represented as mean ± SEM. The significant difference is presented as *P < 0.05, Student’s t test. The level of GAPDH is used as a housekeeping gene to normalize the Western blots data. (A) Effect of DHA on the expression of WAT specific genes (B) Effect of EPA on the expression of WAT specific genes

**Figure (3):**
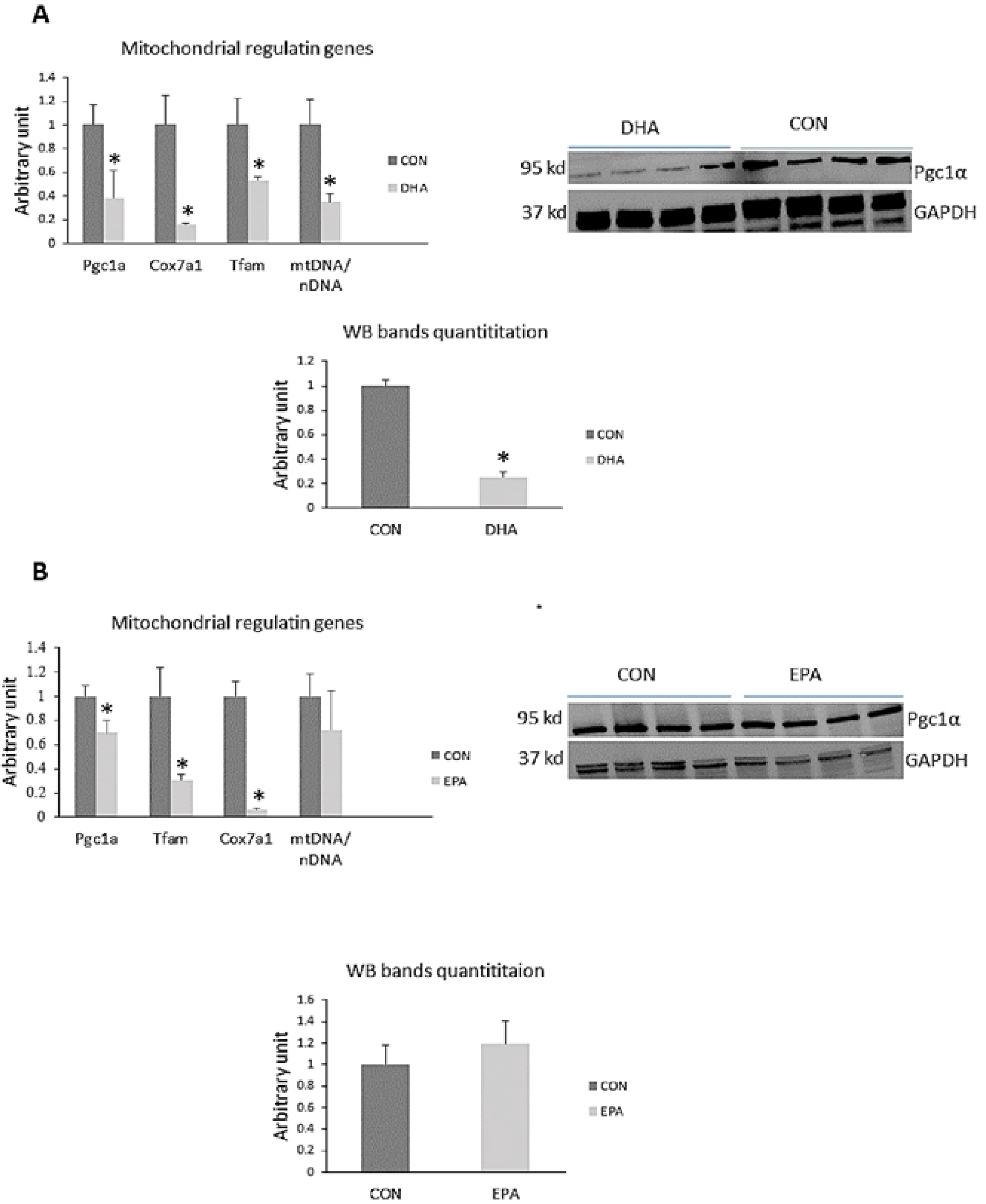
Representative image of western blot, densitometric analysis, and quantitative RT-qPCR analysis of the expression levels of genes regulating mitochondrial biogenesis in C2C12 cells during the trans-differentiation into white adipocytes by a hormonal cocktail. The cells are treated with differentiation induction medium (DIM) in the absence (CON) or presence of individual doses of 50 µM EPA and 50 µM DHA for 10 days. All the values of RT-qPCR are normalized to the expression levels of 18s genes and calculated in arbitrary units, n = 6. Data are represented as mean ± SEM. The significant difference is presented as *P < 0.05, Student’s t test. The level of GAPDH is used as a housekeeping gene to normalize western blots data. **(A)** Effect of DHA on the expression of mitochondrial biogenesis regulating genes. **(B)** Effect of EPA on the expression of mitochondrial biogenesis regulating genes.

#### 3.1.3. DHA reduced the expression of genes regulating mitochondrial biogenesis and mitochondrial DNA (mtDNA) replication

To investigate the effect of EPA and DHA on mitochondrial biogenesis, the expression levels of *PGC1α*, *Cox7α1*, and *Tfam* in C2C12 muscle cells undergoing white adipogenesis differentiation were measured by real-time quantitative PCR, and the ratio of mtDNA to nuclear DNA (nDNA) was determined. mRNA levels of *PGC1α*, *Tfam*, and *Cox7α1* were significantly reduced in DHA treated group (62% ± 3.8, 47% ±2.5, and 84% ± 0.9 respectively) and EPA treated group (30%±10.6, 70%±4.7, and 94% ± 0.8 respectively). Consistently, PGC1α protein content was reduced in DHA treated group but not in EPA treated cells compared to control cells. In alignment, the mtDNA/nDNA ratio decreased by (65 % ±6.6) in the cells treated with 50 μM DHA; however, there were no significant differences for mtDNA copy number in the group treated with 50 μM EPA (**Figure 4).**

**Figure 4:**
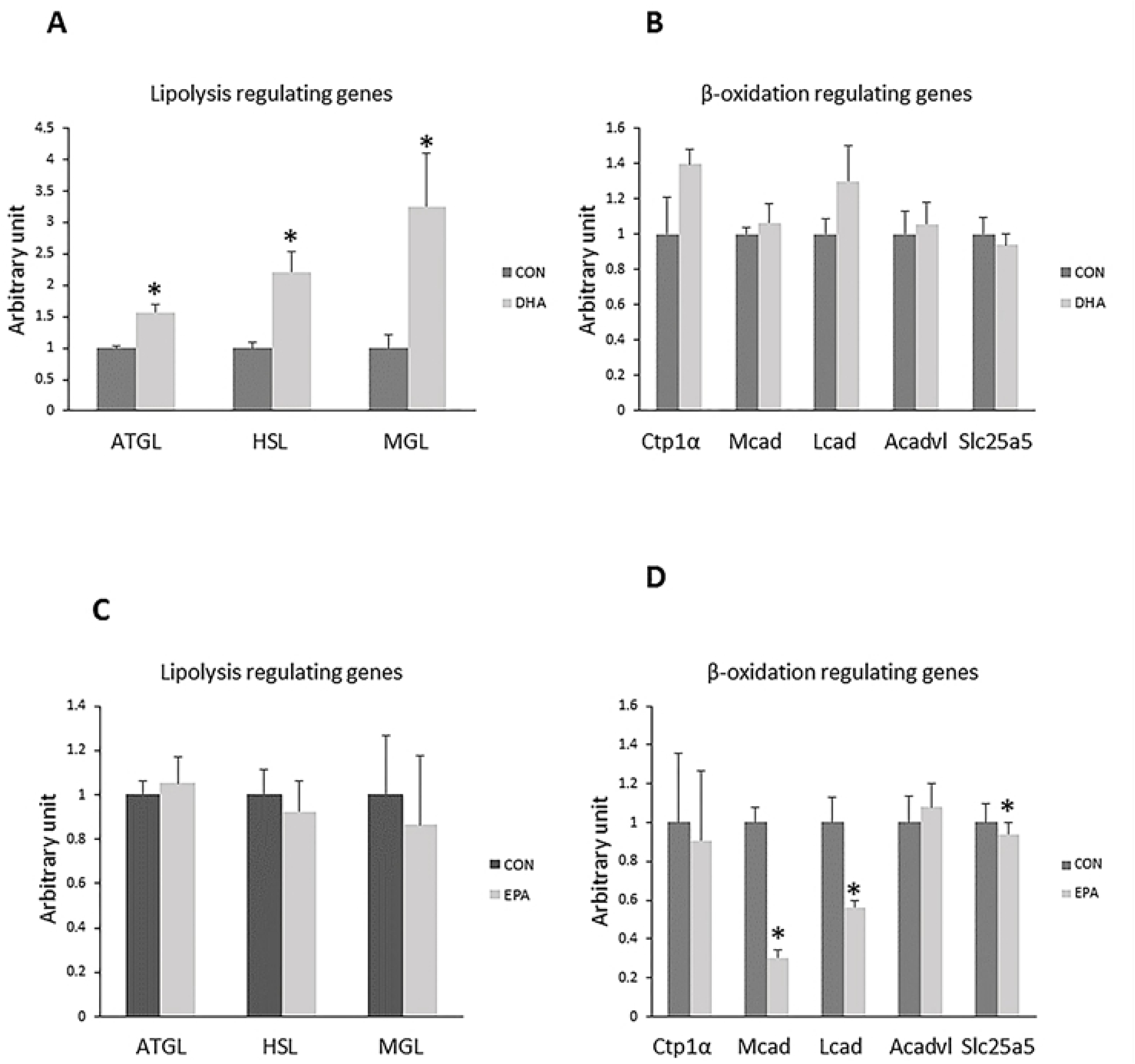
Quantitative RT-qPCR analysis of the expression levels of genes regulating basal lipolytic and β-oxidation activity in C2C12 cells during their reprograming route into white adipocytes by a hormonal cocktail. The cells are treated with differentiation induction medium (DIM) in the absence (CON) or presence of individual doses of 50 µM EPA and 50 µM DHA for 10 days. The values are normalized to the expression levels of 18s genes and calculated in arbitrary units, n = 6 **(A)** Effect of DHA on the expression of genes regulating basal lipolysis. **(B)** Effect of DHA on the expression of genes regulating lipid uptake and β-oxidation **(C)** Effect of EPA on the expression of genes regulating basal lipolysis. **(D)** Effect of EPA on the expression of genes regulating lipid uptake and β-oxidation. Data are represented as mean ± SEM. The significant difference is presented as *P < 0.05, Student’s t test.

#### 3.1.4. Effects of EPA and DHA supplementation on lipolysis and β-oxidation regulating genes

Our data exhibited an increase in basal lipolytic activity in DHA treated group but not in EPA treated group when compared to control. The mRNA levels of *ATGL*, *HSL*, and *MGL* were significantly increased in DHA treated group (56% ± 13.6, 121 % ±32.1, and 225% ± 84.7 respectively); whereas, the mRNA expression of *ATGL*, *HSL*, and *MGL* genes was not altered between EPA treated and control cells. On the other hand, no significant changes between DHA and control groups were observed in the expression levels of genes regulating fatty acids uptake and β-oxidation such as *CPT1α*, *LCAD*, *MCAD*, ACADVL, and SLC25a5. EPA treatment, in contrast, down-regulated the expression levels of *LCAD*, *MCAD*, SLC25a5 by (44% ± 0.4, 70% ± 4, and 49% ± 3.2 respectively), but not the expressions of ACADVL and *CPT1α*. (**Figure 4)**

#### 3.1.5. Effect of EPA and DHA supplementation on lipid droplets formation and morphological changes

Our findings revealed noticeable morphological changes in DHA and EPA treated cells. The absence of multinucleated myofibers and formation of rounded cells was observed in both EPA and DHA treated groups. However, the formation of large monocular bright red lipid droplets was only observed in the DHA treated group. Lipid accumulation in the different groups was confirmed by quantitative measurements of ORO which exhibited a dramatic increase in DHA treated group in comparison to control and EPA treated groups by (76%) (**Figure 5)**. These results were consistent with those obtained from PCR and western blot, implying that C2C12 lost their myogenic capacity once exposed to EPA and/or DHA treatment. However, only DHA had the capability of transdifferentiating C2C12 cells into white-like adipocytes trapping triglyceride in the form of large lipid droplets.

**Figure 5:**
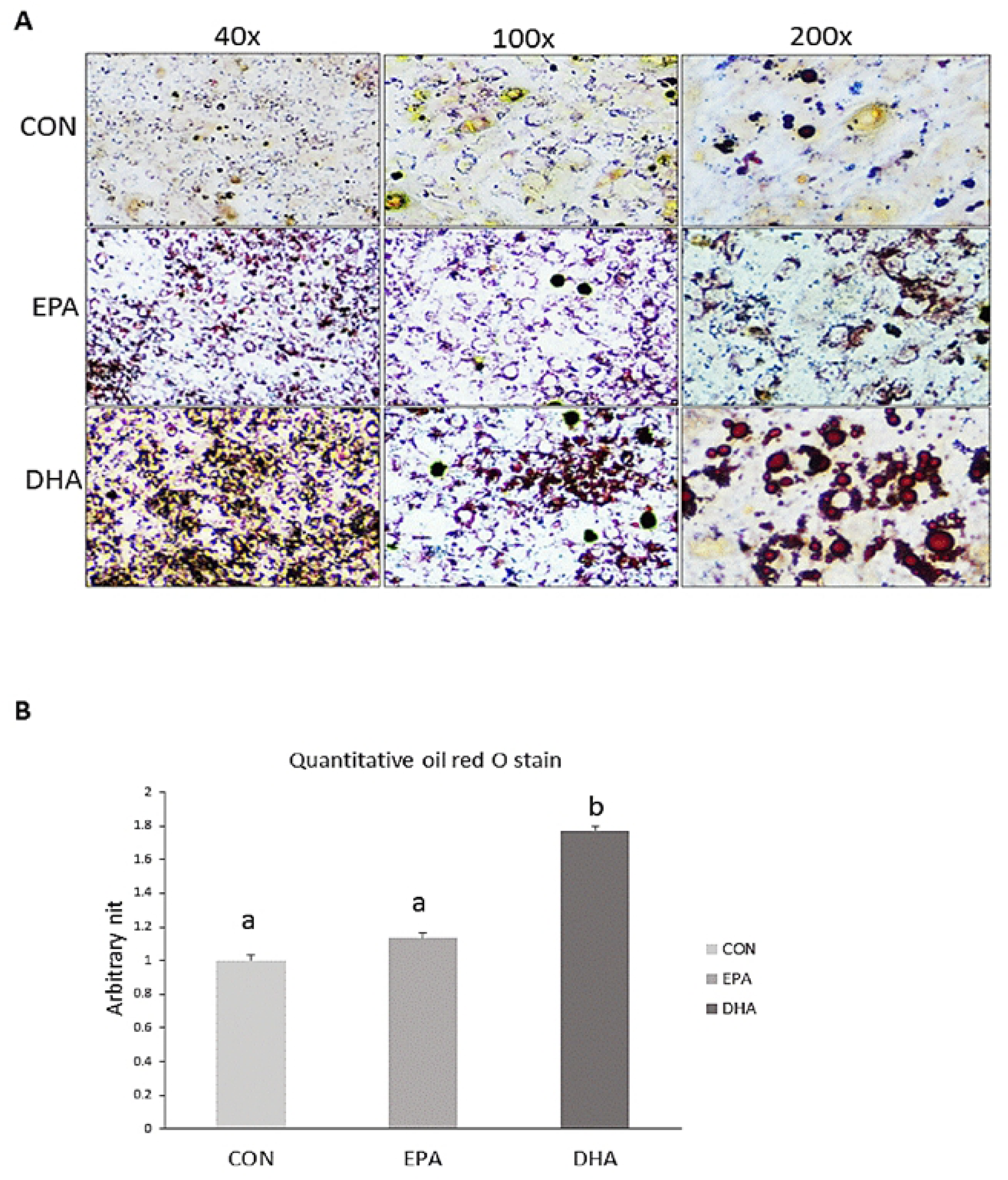
Analysis of morphological changes and lipid droplets formation by Oil Red O (ORO)- staining of C2C12 cells undergoing differentiation into white adipogenesis. The cells are treated with differentiation induction medium (DIM) in the absence (CON) or presence of individual doses of 50 µM EPA and 50 µM DHA for 10 days. **A)** Microscopic view of trans-differentiated cells stained with ORO from different groups. **B)** Quantitative measurement of ORO stain from the different groups. Data is represented as mean ± SEM, b means P < 0.05, ANOVA; n = 4.

### 3.2. Effect of EPA and DHA separately on brown adipogenic differentiation of C2C12 cells

#### 3.2.1. Effect of EPA and DHA supplementation on BAT specific genes

The expression levels of brown adipocytes specific markers including uncoupling protein 1 (*UCP1*, master thermogenic protein) cell death-inducing DFFA-like effector A (*CIDEA*), and PR domain containing 16 (*PRDM16*) was not affected by DHA and EPA treatment. However, genes regulating BAT thermogenesis such as peroxisome proliferator-activated receptor gamma coactivator-1 alpha (*PGC1α*), uncoupling protein 3 (*UCP3*), and Type II iodothyronine deiodinase (*DIO2*) declined in both EPA (68%± 10.0, 93% ± 2.0, and 61% ±2.7, respectively) and DHA treated cells (55%± 10.9, 39 % ± 4.3, and 51 % ± 4.3 respectively) in comparison to control groups. On the other hand, there was no significant difference in PGC1α protein level in EPA treated group when compared to control group **(Figure 6)**. It was apparent that DHA treated group impaired the course of C2C12 cells differentiation into brown adipocytes independent of changing *UCP1* level.

**Figure 6:**
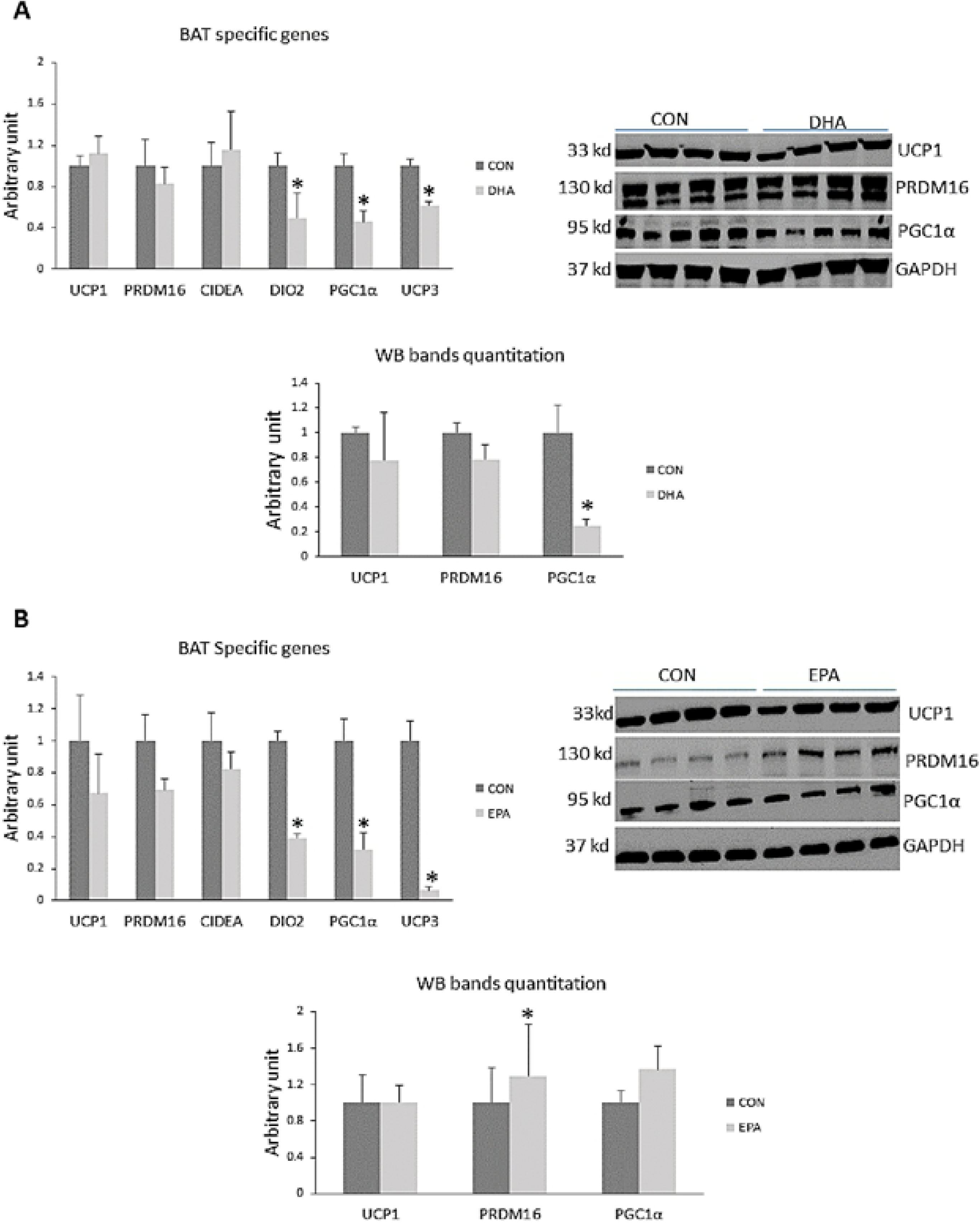
Representative image of western blot, densitometric analysis, and quantitative RT-qPCR analysis of the expression levels of brown adipogenesis master genes in C2C12 cells during the trans-differentiation into brown adipocytes by a hormonal cocktail. The cells are treated with differentiation induction medium (DIM) in the absence (CON) or presence of individual doses of 50 µM EPA and 50 µM DHA for 7 days. All the values of RT-qPCR are normalized to the expression levels of 18s genes and calculated in arbitrary units, n=6. The level of GAPDH is used as a housekeeping gene to normalize the Western blots data **(A)** Effect of DHA on the expression of brown adipogenesis regulating genes. **(B)** Effect of EPA on the expression of BAT signature genes. Data are represented as mean ± SEM. The significant differences were presented as *P < 0.05, Student’s t test.

#### 3.2.2. Effect of treatment of EPA and DHA on genes regulating mitochondrial biogenesis

These muscle cells and BAT are highly enriched with mitochondria, which are responsible for orchestrating the oxidative capacity through the activities of fatty acids oxidation enzymes and respiratory chain components. Thus, reduced mitochondrial density and function impair the thermogenic capacity of cells. We found that the expression levels of *PGC1α* were significantly reduced in groups exposed to treatment with EPA and DHA (68%± 10.0 and 55%± 10.9 respectively); while, *Errα* and *PGC1β* showed no difference between control groups and those treated with DHA or EPA. On the proteomic level, PGC1α was significantly reduced in the DHA treated group only. **(Figure 7)**.

**Figure 7:**
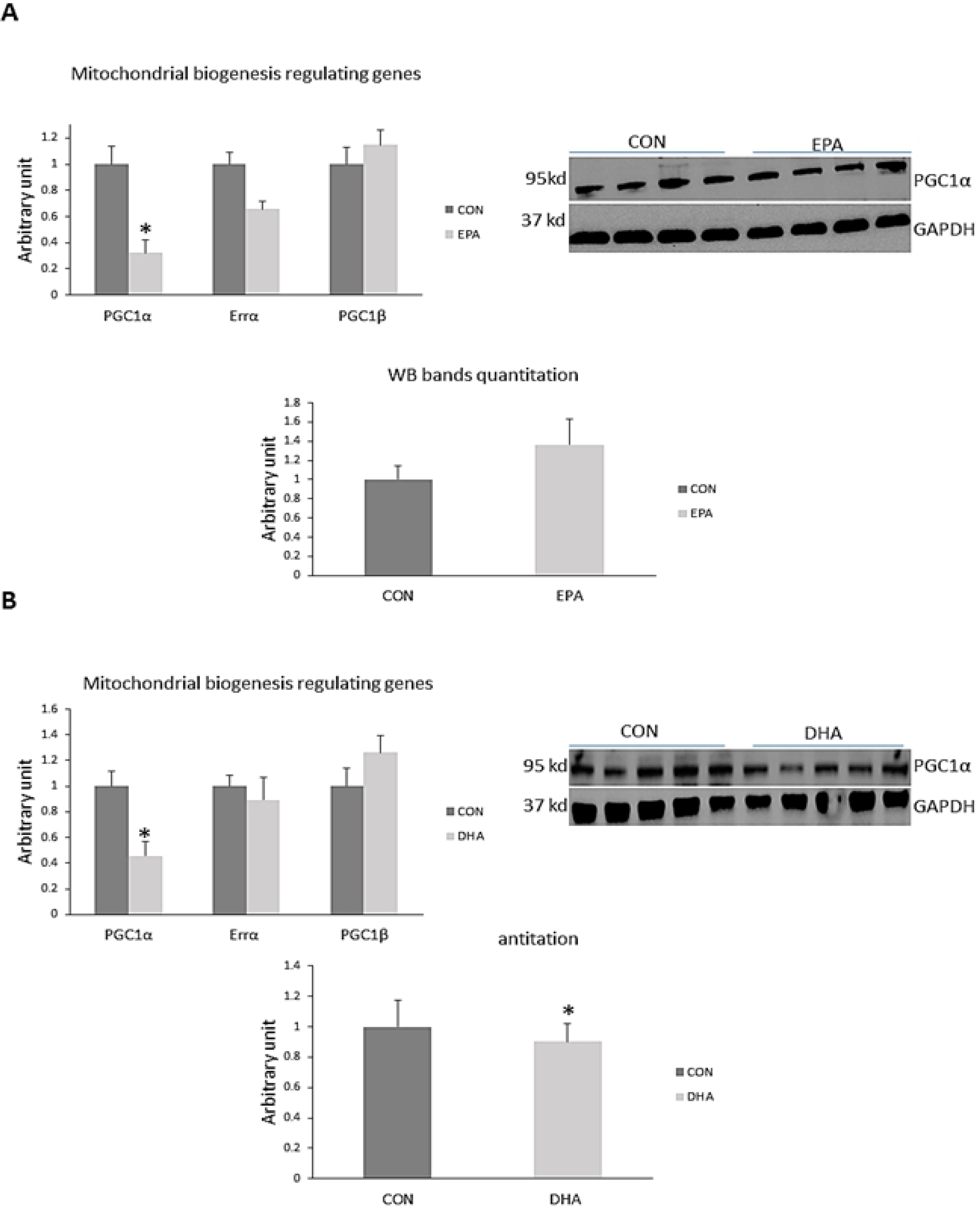
Representative image of western blot, densitometric analysis, and quantitative RT-qPCR analysis of the expression levels of key genes regulating mitochondrial biogenesis in C2C12 cells during the trans-differentiation into brown adipocytes by a hormonal cocktail. The cells are treated with differentiation induction medium (DIM) in the absence (CON) or presence of individual doses of 50 µM EPA and 50 µM DHA for 7 days. All the values of RT-qPCR are normalized to the expression levels of 18s genes and calculated in arbitrary units, n= 6. Data are represented as mean ± SEM. The significant difference is presented as *P < 0.05, Student’s t test. The level of GAPDH is used as a housekeeping gene to normalize the Western blots data **(A)** Effect of DHA on the expression of mitochondrial biogenesis regulating genes. **(B)** Effect of EPA on the expression of mitochondrial biogenesis regulating genes.

#### 3.2.3. Effect of EPA and DHA treatment on genes regulating ETC work

Co-factors produced from TCA and β-oxidation are directed to the electron transport chain (ETC) to be used in uncoupling respiration to produce heat which is the main function of brown adipocytes. We found that the expression levels of *COX7α1*, *COX8β* were by far down-regulated in EPA by (61 % ± 6.3 and 87% ± 2.6 respectively) and DHA treated groups (68 % ± 7.2 and 97 % ± 2.6 respectively) when compared to control groups; whereas, the mRNA expressions of *COX5*, and *ATP5jα* were no significantly affected by treatments with EPA or DHA. Also, non-significant changes were observed in protein levels of complex II and complex III in both groups exposed to EPA and DHA treatment upon comparison to control groups. (**Figure 8).**

**Figure 8:**
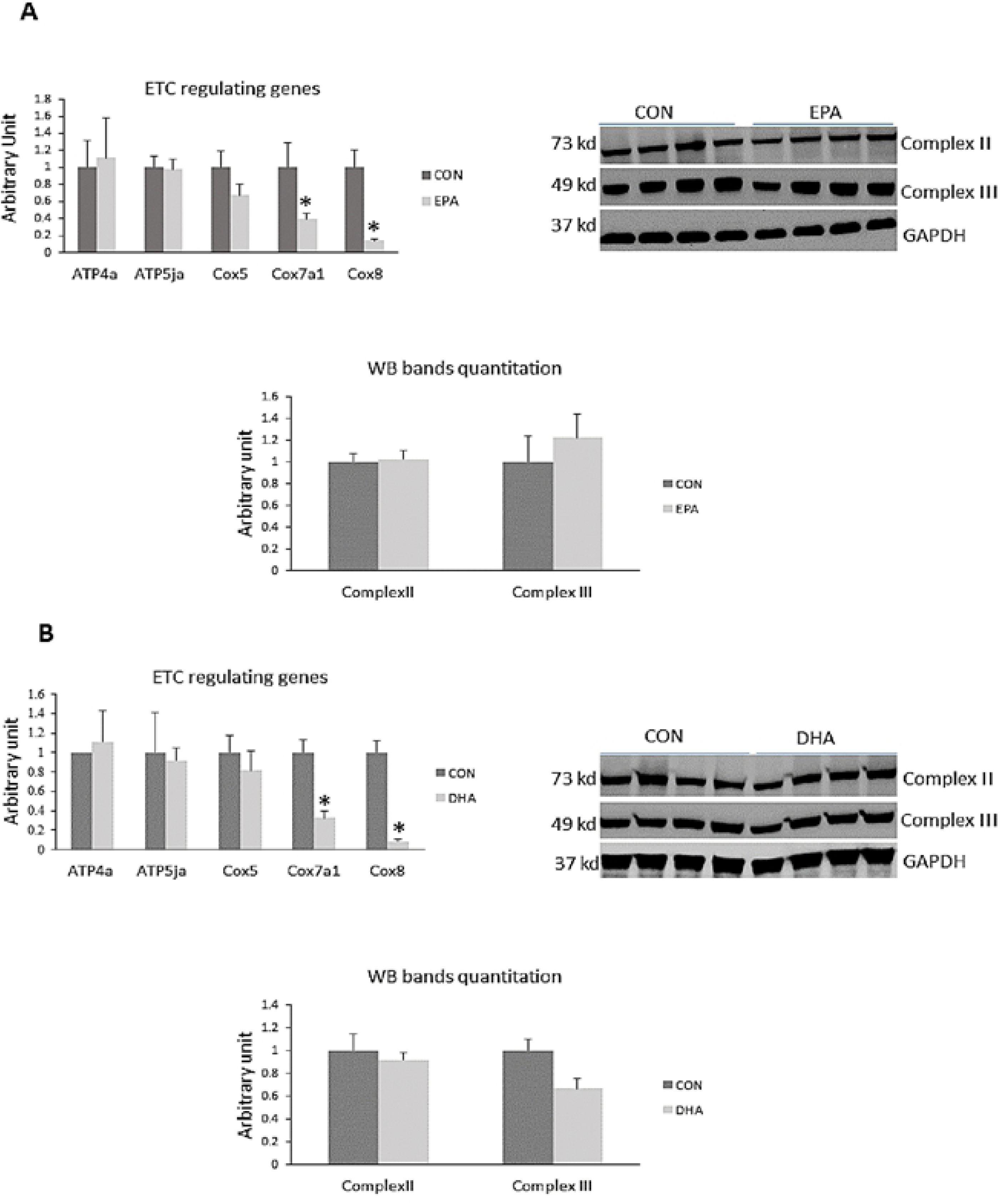
Representative image of western blot, densitometric analysis, and quantitative RT-qPCR analysis of the expression levels of some electron transport chain regulating genes in C2C12 cells undergoing differentiation into brown adipocytes by a hormonal cocktail. The cells are treated with differentiation induction medium (DIM) in the absence (CON) or presence of individual doses of 50 µM EPA and 50 µM DHA for 7 days. All the values of RT-qPCR are normalized to the expression levels of 18s genes and calculated in arbitrary units, n= 6. Data are represented as mean ± SEM. The significant difference is presented as *P < 0.05, Student’s t test. The level of GAPDH is used as a housekeeping gene to normalize the Western blots data. **(A)** Effect of DHA on the expression of ETC regulating genes. **(B)** Effect of EPA on the expression of ETC regulating genes.

#### 3.2.4. Effect of EPA and DHA supplementation on lipid accumulation and morphological changes

Our findings showed notable losses of multinucleated myotubes and changing the morphology of the cells into rounded cells filled with lipid droplets (**Figure 9)**. Also, an increase in lipid droplets size and number was observed in DHA treated cells when compared to other groups **(SF1, ST1)**. These data were consistent with the results obtained from the quantitative measurement of ORO which exhibited a dramatic increase in DHA treated group when compared to EPA and control groups by (200 % ±27.3) (**Figure 9)**.

**Figure 9:**
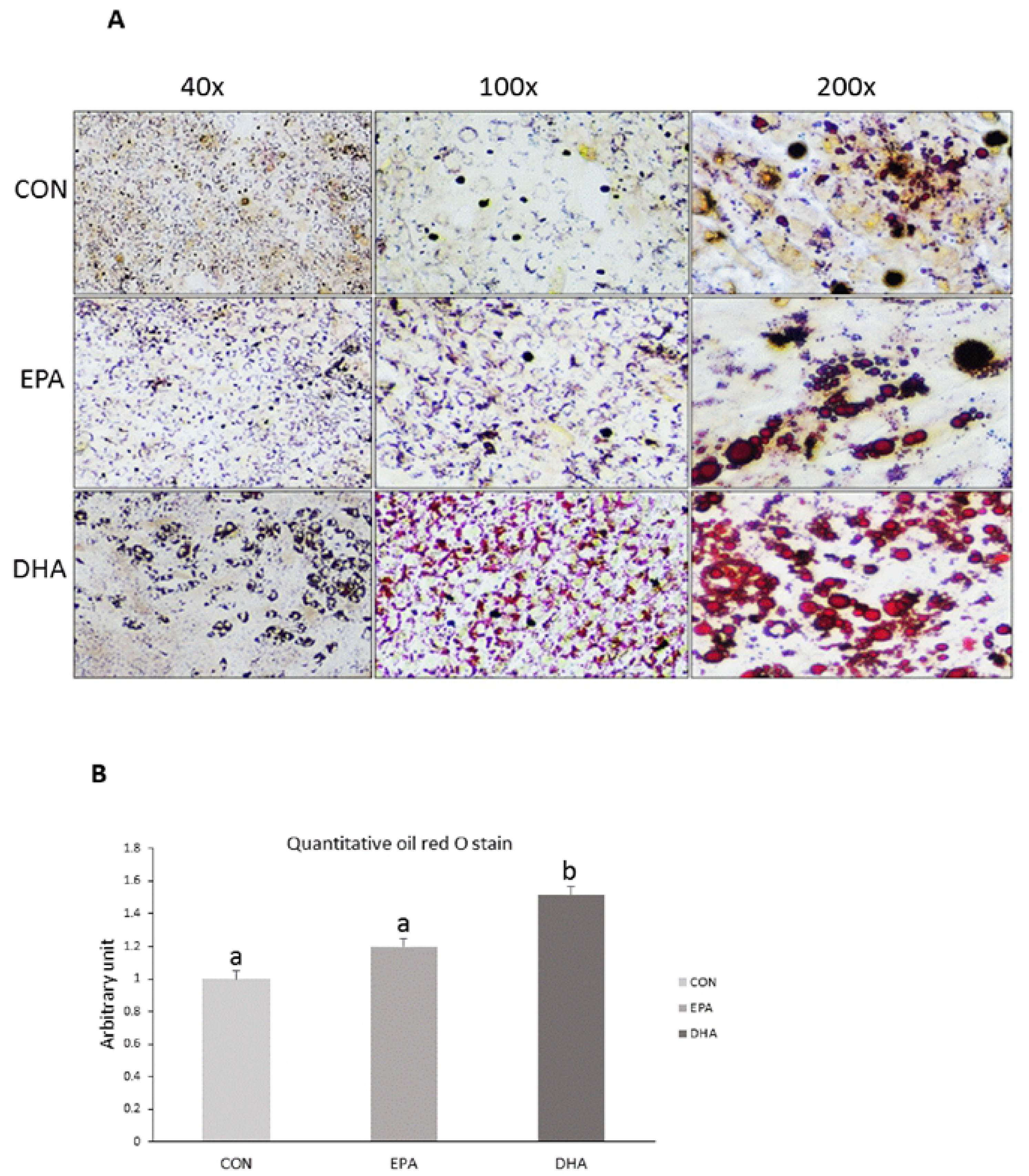
The effect of EPA and DHA separately on lipid accumulation in C2C12 myoblasts after induction of brown adipogenesis. The cells are treated with differentiation induction medium (DIM) in the absence (CON) or presence of individual doses of 50 µM EPA and 50 µM DHA for 7 days. (**A**) Representative images of ORO staining of cells from different groups exhibiting morphological changes and lipid droplet formation (**B)** Quantitative measurement of ORO staining in control and fatty treated groups. Significant difference between the two groups are measured in arbitrary unit and data are represented as mean ± SEM. Letters represents significant difference where b means P < 0.05, ANOVA; n=4.

**Figure 10:**
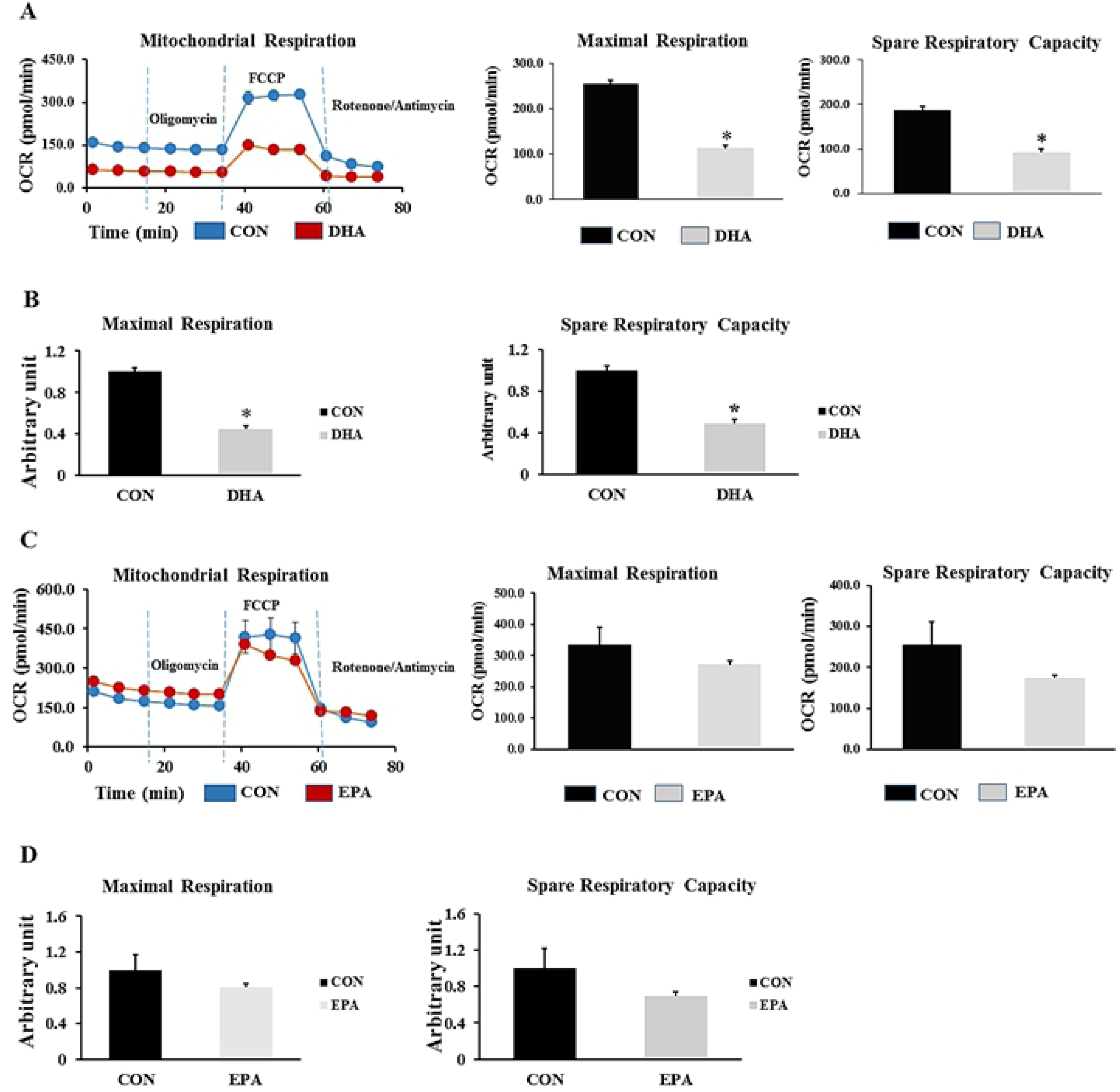
The effect of EPA and DHA separately on metabolic rate of C2C12 cells undergoing differentiation into brown adipogenesis. The cells are treated with differentiation induction medium (DIM) in the absence (CON) or presence of individual doses of 50 µM EPA and 50 µM DHA for 7days. Vertical dashed lines indicate the times of addition of oligomycin (2 µM), FCCP (0.7 μM), and antimycin A (10 µM) **(A)** the effect of DHA treatment on maximal respiration and spare respiratory capacity. OCR traces are expressed as pmol O2 per min in C2C12 cells and normalized to protein concentration **(B).** The Relative differences in maximal respiration and spare respiratory capacity of control and DHA treated group **(C)** The effect of EPA supplementation on maximal respiration and spare respiratory capacity. OCR traces are expressed as pmol O2 per min in C2C12 cells and **(D)** the relative changes in maximal respiration and spare respiratory capacity are measured in arbitrary units, n=3 and 12 measurements. Data are represented as mean ± SEM. The significant difference is presented as *P < 0.05, Student’s t test.

#### 3.2.5. DHA but not EPA reduced mitochondrial respiration and spare capacity

The maximal mitochondrial respiration and spare capacity were suppressed in DHA treated cells by (56%± 3.0 and 51% ± 4.1 respectively). On the other hand, no differences were found between control and EPA treated groups. Taken together, the reduction of mitochondrial content and metabolic efficiency due to DHA might be attributed to suppression of the mitochondrial content resulted from suppressing of PGC1α expression **(Figure10).**

## 4. Discussion

In addition to its contribution to more than 40% of body mass, skeletal muscle is considered a major site of glucose disposal because of its enrichment with insulin receptors. In this sense, it was reported that roughly 80% of circulating glucose is used for fuel daily muscular activities (30). A positive correlation has been revealed between intramuscular lipid infiltration and lipotoxicity and related insulin resistance in humans as muscle tissue is not as efficient as adipose tissue in lipid storage (31, 32). Reprogramming of skeletal muscle precursor cells into functional adipocyte only occurs after stimulation by certain drugs and cytokines or concurrently with pathophysiological conditions (17, 33–36). Only muscle precursors and satellite cells can be induced to Trans-differentiate into adipocyte in response to external stimuli. However, the conversion of myotubes at the end stage of differentiation into adipocytes is not likely feasible (34). Generation of fat cells from myogenic precursors is a complicated process which requires suppression of myogenic genes (37). Here, we identified that both EPA and DHA have the ability to inhibit myotube formation in vitro. However, DHA but not EPA could effectively induce trans-differentiation of myoblasts into white-like adipocytes that might subsequently lead to the accumulation of intramyocellular lipid. In our study, DHA enhanced the trans-differentiation of myoblasts in to white -like adipocytes, which could be due to several possible mechanisms: a) inhibiting the terminal differentiation of myoblasts into mature myotubes via down-regulating the expression of myogenic marker genes such as *MyoD1*, *MyoG*, and *MRF4*; b) inducing the expression of white adipogenic gene including *PPARγ*, *C/EBPβ*, *C/EBPα*, *Ap2*, *FAT*, *Bmp4*, and *Zfp423*; and c) impairing the acquisition of functional brown adipocyte phenotype through reducing the function and density of mitochondria leading to lipid accretion instead of utilization through oxidation.

The first evidence of DHA induced C2C12 cells reprogramming into white adipocyte like phenotype is the ectopic expression of adipogenesis key regulators, *PPARγ*, *C/EBPβ*, and *C/EBPα* that drives C2C12 to loss their myogenic identity and switches into the adipocytes like lineage. Inhibition of myotubes formation can be attributed to reducing the expression of myogenesis regulating genes particularly *MyoD1*, *MyoG*, and *MRF4*. However, it was reported that muscle progenitor cells could restore their myogenic capacity once *PPARγ* is disconnected from its activator, indicating that the trans-differentiation process is temporary and depends on the sustained expression of *PPARγ* and *C/EBPα* (17). It has been demonstrated that *PPARγ* is the key regulator of adipocyte formation from non-adipogenic precursors, where its expression is sufficient and indispensable for the terminal differential of fibroblast into mature adipocytes (16, 38). A comparative study was conducted to elucidate the necessity of the ectopic expression of *C/EBPα* and *PPARγ* in committing mouse embryonic fibroblasts (MEFs) into adipocyte lineage. They found that the *PPARγ* ectopic expression is sufficient to induce the trans-differentiation of *C/EBPα*/^-^ MEFs into adipocytes in vitro (39); whereas, enforcing the expression of *C/EBPα* in *PPARγ*/^-^ MEFs was not adequate to induce the reprogramming process (40). These results indicated that C/EBPα is not obligate for adipocyte differentiation, while PPAR ectopic expression is indispensable for driving non-adipogenic progenitors into adipocyte lineage. The myogenesis process is regulated by a group of important MRFs, including *Myf5*, *Myod1*, MyoG and Myf6 (*Mrf4*) (41). It has been demonstrated that Myf5+ progenitors have the potential to switch between two different lineages including myoblasts and brown preadipocytes in response to surrounding environment. Persistent expression of *MyoD1* drives cell commitment toward the myogenic fate while commitment to brown adipocytes requires the suppression of *MyoD1* (42). However, suppression of *MyoD1* and related trans-differentiation of myogenic precursors into adipocytes is not accompanied with *UCP1* expression, and its expression required the interaction of other factors such as Notch signaling and sympathetic innervation to be induced (43, 44). These observations are in agreement with our results suggesting that white like adipocytes can be arisen from Myf5+ myoblasts upon their exposure to *PPARγ* activators. The second evidence is that the DHA treated group showed increased expression level of white adipocytes specific genes including *FAT*, *Bmp4*, and *Zfp423*. It was reported that Bmp4 promotes brown adipocytes switching into white adipocyte phenotype by down-regulation the expression of brown adipogenesis specific markers and up-regulation of white adipogenesis signature genes. Also, it suppressed lipolysis and inhibited oxygen consumption rate (OCR) followed by reducing the thermogenic capacity of cells (45). *Zfp423* was reported to exert a profound effect on adipogenesis regulation and associated intramuscular lipid overload in beef cattle. Stromal vascular cells isolated from beef samples were induced to Trans-differentiate into adipocytes via enforcing the expression of *Zfp423*, which further induced the expression of adipogenesis key regulators, *PPARγ* and *C/EBPα* (46). Additionally, *Zfp423* suppressed the thermogenic capacity of brown adipocytes through reducing the activity of Ebf2 leading to blunting the expression of *Prdm16*, a key transcription factor required for brown adipogenesis (47). Moreover, reprogramming white adipocytes into beige-like adipocytes requires inactivation of *Zfp423*, which is highlighted as a therapeutic option to counter obesity (48). Consistently, we found that EPA supplementation reduced the expression levels of these two white adipocytes specific markers (*Bmp4* and *Zfp423*), referring to its inverse association to the expression of transcription factors enhancing white adipose tissue expansion. Taken together, DHA supplementation was associated up-regulation of whitening signature genes induced C2C12 cells to trans-differentiate into white adipocyte like phenotype; whereas, inhibiting the expression and protein levels of myogenesis regulating genes may prevent myotube formation but not necessarily enhance the trans-differentiation of myoblasts into adipocyte lineage which was observed in the cells exposed to EPA. The commitment of C2C12 cells into white-like adipocytes appears to be entirely mediated by *PPARγ* overexpression while activating the Bmp4 - Zfp423- PPARγ pathway favors promoting the white adipogenesis while prevents the development of brown adipocytes phenotype (47).

The third evidence of C2C12 cells reprogramming into white like adipocytes upon DHA exposure is down-regulation the expression of genes responsible for mitochondrial thermogenic capacity including *PGC1α*, *Cox71α*, *Cox8β*, *Dio2*, and *UCP3*. *PGC1α* was identified as critical elements in creating brown adipose tissue and enhancing their function through stimulating mitochondrial biogenesis and driving *UCP1* expression by stimulating its promoter (49). Down-regulation of *PGC1α* expression in C2C12 cells exposed to EPA and DHA is consistent with other studies in which the level of PGC1α was considerably decreased in mice treated with n-3 fatty acids alone or in combination with dexamethasone (50). However, our findings showed non-significant change in *UCP1* gene expression and protein level despite of inhibiting the expression of its activator PGC1α indicating the potential regulation of *UCP1* production by another molecular pathways. Also, our results showed that EPA treatment has a massive effect on *PGC1α* mRNA expression but not at the level of protein in both experiments. Furthermore, reducing the expression level of *Dio2*, a well-known regulator of thermogenesis process, could be highlighted as another indicator of myoblast conversion into white like adipocytes phenotype in this study. Type II iodothyronine deiodinase (Dio2) induced thyroxin (T4) conversion into its active form triiodothyronine (T3) is the rate-limiting step in stimulating the thermogenic capacity of cells. Cold exposure and norepinephrine associated increased thermogenesis can be attributed to up-regulation of Dio2 (51). Therefore, down-regulating the expression of Dio2 may reduce thermogenesis. Consistently, Mehus and Picklo (52) revealed a negative correlation between EPA and DHA supplementation and Dio2 expression level in mice cerebral cortex. However, unlike in DHA treated group, EPA treatment induced down-regulation of thermogenic regulating genes was not accompanied by mitochondrial dysfunction as no change was observed in maximal respiration and spare respiratory capacity. Also, ORO staining showed no change in lipid droplets size and number.

The fourth evidence of C2C212 conversion into white-like adipocytes after DHA treatment is reducing the expression of genes regulating mitochondrial biogenesis in the DHA treated group. Given that the phenotype of adipocyte cells can be identified based on their mitochondrial content, we investigated the effect of DHA and EPA supplementation on mitochondrial density and its association with mitochondrial function manifested by maximal respiration and spare respiratory capacity. Accordingly, we measured the relative expression and protein levels of mitochondrial biogenesis regulating key genes. While DHA reduced the gene expression and protein level of PGC1α and Tfam, the expression of other genes involved in regulating mitochondrial biogenesis such as *PGC1β* and *Errα* were not changed. It was reported that *PGC1α*, but not *PGC1β* is critical for orchestrating mitochondrial thermogenic capacity. However, the biogenesis process can be comprised by reducing the expression of either of them (53). Additionally, It was demonstrated that *PGC1α* regulates the expression of *NRF1* and *NRF 2* which are nuclear transcription factors responsible for regulating the expression of mitochondrial transcription factor A (*Tfam*), controlling mitochondrial DNA content inside the cells (37, 54). DHA-induced reducing mitochondrial copy number in C2C12 cells, during the course of differentiation into white adipocytes, was predicted from decreased mtDNA/nDNA ratio. Reducing mitochondrial content in the DHA treated group can be attributed to suppressing PGC1α protein level and associated decreased Tfam expression. Down-regulating Tfam expression could be followed by respiratory chain disorder and mitochondrial dysfunction. On the other hand, the suppressing effect of EPA on *PGC1α* in both experiment was only noticed on the transcriptome level while protein production was unchanged when compared to control group and was concurrent with normal mitochondrial function. It is to be included that the DHA associated mitochondrial dysfunction observed in this study may be attributed to reducing the density of mitochondria. Moreover, the observed reduction in mitochondrial maximal respiration and spare respiratory capacity in group treated with DHA and BIDM can be linked to its incorporation in the mitochondrial membrane. Correspondingly, it was stressed that DHA induced structural changes in lipid components of the inner mitochondrial membrane isolated from mice cardiocytes led to reducing the enzymatic activities of OXPHOS subunits including complexes I, IV, V, and I+III. In consistent with our study, the findings exhibited suppressing the activity but not the expression level of OXPHOS regulating subunits (55). Additionally, it has been recently discovered that DHA is an essential factor in remodeling mitochondrial phospholipidome (56–58). Given that both EPA and DHA has the same effect on the set of genes regulating mitochondrial content and thermogenesis but only DHA treated group exhibited insufficiency in mitochondrial work, we think that DHA mediated suppressing mitochondrial biogenesis as a result of reducing PGC1α protein production is the main reason behind mitochondrial dysfunction and related impairing the acquisition of brown adipocyte like phenotype. However, the hypothesis of linking DHA incorporation into cell membrane to the generation of brown adipocytes with less mitochondrial content and function is still feasible as it has been mentioned that DHA is the key player in remodeling the biochemical structure of cell membrane mainly lipid rafts; whereas, EPA is mainly implicated in orchestrating other central processes including immunity and inflammation (59, 60). Another reason might be linked to reducing the function of mitochondria in DHA treated groups is the potential excessive production of mitochondrial ROS in MAPKs activation dependent manner (61). They found that using antioxidant reversed DHA-induced mitochondrial malfunction and enabled restoring the mitochondrial respiration rate.

The fifth evidence was the observed increased in lipid droplets size and number in DHA treated group. It is well known that one of the main metabolic characteristics of adipocytes is intracellular lipid storage, where white adipocyte cells are characterized by their monocular lipid droplets while brown adipocytes are enriched with small and multillocular droplets. In this study, we found that in both experiments there was an increase in the number and size of lipid droplets in DHA treated group in comparison to control and EPA treated groups. In agreement, a positive correlation between EPA and DHA supplementation and intracellular lipid accumulation was observed. A study emphasized that omega-3 PUFA associated increase in intramyocellular deposition of lipid droplets can be attributed to TGA synthesis (62). Additionally, DHA related inhibition of thermogenesis and up-regulation of WAT regulating genes such as PPARγ, *C/EBPα*, *FAT*, *BMP4*, and *ZFP423* observed in this study may be the reason behind increasing lipid droplets size and number.

The sixth evidence is DHA induced lipid re-esterification. Our findings revealed up-regulation of genes regulating basal lipolysis in group treated with combination of DHA and WIDM. In line with that, the results exhibited an increase in lipid trapping inside the cells and unchanging the levels of set of genes known to be involved in enhancing fatty acid uptake and catabolism. Thus, we can conclude that free fatty acids resulted from triglyceride breakdown are re-uptake and re-esterified by the cells again. It was reported that lipid handling through several interconnected process such as lipolysis, glyceroneogenesis, and FA re-esterification is one of the main features of white adipose tissue (63). On the other hand, we found that EPA induced suppressing of some genes regulating β-oxidation. We think lacking β-oxidation capacity in EPA treated group may not be linked to mitochondrial insufficiency, particularly the results did not exhibit changing maximal respiration and spare respiratory capacity; instead, metabolic flexibility allows cells to easily switch between glucose and lipid oxidation. In line with that, the non-mitochondrial respiration was enhanced in EPA treated group when compared to control (**SF2**) indicating the preferential role of EPA in enhancing the capacity of muscle cells in substrates handling (64).

## 5. Conclusion

We can conclude that C2C12 cells losses their myogenic capacity once exposed to EPA and DHA treatment. DHA is an adipogenic factor inducing C2C12 cells trans-differentiate into white-like adipocytes by increasing the expression level of its ligand *PPARγ*, the master regulator of adipogenesis process. *PPARγ* activation accompanied with up-regulation of whitening signature genes like *C/EBPα*, *C/EBPβ*, *AP2*, *BMP4*, and *Zfp423*. Also, adipocytes derived from C2C12 treated with DHA decreased thermogenic capacity and reduced mitochondrial copy number and activity. Mitochondrial dysfunction observed in DHA treated group might be attributed to inhibition of genes and proteins known to be implicated in regulating mitochondrial biogenesis and thermogenic capacity such as *PGC1α*, *Dio2*, *UCP3*, and *Tfam*. The results showed no change in *UCP1* level between groups indicating that the effect of DHA on brown adipogenic differentiation of C2C12 cells is independent of changing *UCP1*expression. Adipocytes arise from C2C12 after DHA treatment were enriched with big uniocular lipid droplets. The obvious increase in size and number of lipid droplets in DHA treated group may have been caused by incorporation of DHA in the cellular membrane or enhancement of triglyceride synthesis (TGA). Moreover, DHA enhances basal lipolysis and fatty acid re-esterification. EPA is less potent than DHA as adipogenic agent inhibiting the myogenesis process and inversely affecting the expansion of white adipose tissue by reducing the expression of *Bmp4* and *Zfp423*, an important transcriptional factors implicated in regulating white adipogenesis.

## ACKNOWLEDGMENTS

This work was supported by Ministry of Higher Education and Scientific Research - Iraq. The authors would like to thank Dr.Rajaraham for providing Seahorse XFP machine.

## List of abbreviations

N-3-PUFA: n-3 polyunsaturated fatty acids
PPARγ: Peroxisome proliferator-activated receptor gamma
IBMX: 3-isobutyl-1-methylxanthine
WIDM: White adipogenesis differentiation induction medium
BDIM: Brown adipogenesis differentiation induction medium
ORO: Oil Red O stain
UQCRC: Ubiquinol Cytochrome c Reductase
SDHA: Succinate dehydrogenase complex flavoprotein subunit A
18s: 18S ribosomal RNA
C/EBPβ: C/EBP: CCAAT-enhancer-binding proteins beta sub-unit
C/EBPα: CCAAT-enhancer-binding proteins alpha sub-unit
Myf5: Myogenic factor 5
MyoG: Myogenin
AP2: Adipocyte Protein 2
FAT: 
Zfp423: Zinc finger protein 423
Bmp4: Bone morphogenetic protein 4
mtDNA: Mitochondrial DNA
PGC1α: Peroxisome proliferator-activated receptor gamma coactivator 1-alpha
Cox7α1: Cytochrome c oxidase polypeptide 7A1
Cox8β: Cytochrome c oxidase subunit 8 beta
Tfam: Mitochondrial transcription factor A
ATGL: Adipose triglyceride lipase
HSL: Hormone-sensitive lipase
MGL: Monoacylglycerol lipase
CPT1α: carnitine acyltransferase I alpha sub-unit
LCAD: acyl-CoA dehydrogenase, long chain
MCAD: medium-chain acyl-CoA dehydrogenase
ACADVL: very long-chain specific acyl-CoA dehydrogenase
SLC25a5: Solute carrier family 22 member 5
COX5: Cytochrome *c* oxidase subunit 5
ATP5jα: ATP synthase-coupling factor 6 sub unit alpha
ATP4α: ATPase subunit alpha-4
TGF-β: Transforming Growth Factor-β
OXPHOS: oxidative phosphorylation

